# H_2_S is a potential universal reducing agent for Prx6-type peroxiredoxins

**DOI:** 10.1101/2025.02.13.638076

**Authors:** Lukas Lang, Laura Leiskau, Lea Thullen, Marcel Deponte

## Abstract

The absence of a universal reducing agent distinguishes the Prx6-type subfamily of peroxiredoxins from the structurally similar Prx1-type subfamily. A likely explanation for the lack of reactivity of Prx6-type enzymes with common reducing agents is that a Prx6-specific histidyl residue at the bottom of the active-site pocket traps the oxidized enzyme with a potentially hypervalent cysteinyl sulfur atom in an inaccessible fully-folded protein conformation. Here, we analyzed the reduction of oxidized PfPrx6 from the human malaria parasite *Plasmodium falciparum* and human PrxVI by the hydrosulfide ion, HS^−^, as the smallest possible sulfur-containing universal electron donor using stopped-flow kinetic measurements. We show that HS^−^ rapidly reacts with oxidized wild-type PfPrx6 or human PrxVI (but not the histidyl mutant PfPrx6^H39Y^) with a second-order rate constant of >10^8^ M^−1^s^−1^ at pH 7.4. The potential protein-persulfide species of PfPrx6 is neither reduced by thioredoxin nor glutaredoxin and glutathione, but further reacts with an excess of HS^−^ with a second-order rate constant of 6.3×10^3^ M^−1^s^−1^, yielding the reduced enzyme. In summary, we identified HS^−^ as a highly reactive, potential universal electron donor for Prx6-type enzymes in eukaryotes, bacteria, and archaea. Our study marks the starting point for the characterization of the complex reduction pathway of Prx6-type enzymes with potential implications for H_2_S detoxification, redox signaling, and iron-sulfur metabolism.

## Introduction

The identification of a universal physiological reducing agent for oxidized Prx6-type peroxiredoxins is an ongoing quest since the reported characterization of recombinant human PrxVI (hPrxVI) in 1998.^1^ Analogous to other peroxiredoxin subfamilies (EC 1.11.1.15), Prx6-type enzymes are peroxidases that reduce various hydroperoxides and peroxynitrous acid, yielding an enzyme sulfenic acid.^2-8^ However, despite high structural similarity to Prx1-type enzymes,^9^ Prx6-type enzymes have a subfamily-specific active-site histidyl residue,^10-12^ and many oxidized Prx6-type enzymes cannot be reduced by thioredoxins (Trx), glutaredoxins (Grx), or glutathione (GSH).^1,2,4-8,10,12,13^ It is therefore unclear, how oxidized Prx6-type enzymes are reduced again to complete the catalytic cycle. A few exceptions include hPrxVI, which is reduced by glutathione in the presence of glutathione transferase P1-1,^14^ and its yeast homologue, which is directly or indirectly connected to the mitochondrial glutathione pool.^15^ Successful reductions of oxidized Prx6-type enzymes were usually detected for non-physiological dithiothreitol (DTT) in contrast to other low-molecular-weight thiols.^1-7,10,12^ Ascorbate was also shown to reduce some oxidized Prx6 homologues *in vitro*,^16,17^ and H_2_Se derived from selenocysteine was recently reported to form a thioperselenide with hPrxVI.^18,19^ However, Prx6-type enzymes are found in eukaryotes, bacteria, and archaea,^9^ many of which have no corresponding glutathione transferase, or ascorbate or selenocysteine metabolism. Accordingly, several Prx6-type enzymes, including PfPrx6 from the human malaria parasite *Plasmodium falciparum*, showed no activity with a glutathione transferase or ascorbate.^1,2,4,6,12,13^ Here, we analyzed the reaction kinetics between oxidized PfPrx6 or hPrxVI and the H_2_S-derived hydrosulfide ion, HS^−^, as a potential universal electron donor for oxidized Prx6-type enzymes.

## Results and Discussion

### Oxidized PfPrx6 and hPrxVI react rapidly with HS^−^

We previously showed that oxidized PfPrx6 most likely stays in a trapped, fully-folded enzyme conformation involving an interaction between the sulfenic-acid form of the peroxidatic cysteinyl residue and the Prx6-specific histidyl residue at the bottom of the active-site pocket.^8,10,12^ These findings were in accordance with crystal structures of oxidized hPrxVI and ApPrx6 from the archaeon *Aeropyrum pernix*, revealing either a hydrogen bond or a hypervalent sulfur species that involves the active-site cysteinyl and histidyl residue (Fig. 1A).^20-22^ Since oxidized PfPrx6 could only be reduced by DTT but not by other small thiols or ascorbate,^12^ we decided to test HS^−^ as the smallest possible sulfur-containing universal electron donor for inaccessible, fully-folded Prx6-type enzymes using stopped-flow kinetic measurements (Fig. 1A,B). Since the p*K*_a_ values of H_2_S are 7.0 and >14, freshly dissolved Na_2_S can be used as a HS^−^ source. Mixing oxidized wild-type PfPrx6 (PfPrx6^WT^) in the first syringe with variable concentrations of Na_2_S in the second syringe resulted in up to three distinct reaction phases (Fig. 1B). Similar results were obtained for recombinant hPrxVI (Fig. 1C). An extremely rapid Na_2_S concentration-dependent decrease of tryptophan fluorescence was observed within 20-30 ms (Fig. 1B), resulting in a second-order rate constant of (2.8±0.2)×10^8^ M^−1^s^−1^ for PfPrx6^WT^ at pH 7.4 (Fig. 1D, Table 1), which corresponds to a pH-independent value of 3.9×10^8^ M^−1^s^−1^ (based on the HS^−^ protonation state). The first phase occurred during the dead-time for Na_2_S concentrations ≥ 10 μM and was even faster with a second-order rate constant of (5.0±1.2)×10^8^ M^−1^s^−1^ for hPrxVI (Fig. 1B-D, Table 1). A subsequent increase of fluorescence did not depend on the Na_2_S concentration, yielding a rate constant around 11 s^−1^ for PfPrx6^WT^ or 30 s^−1^ for hPrxVI. A third phase was observed at higher Na_2_S concentrations, resulting in an increase of tryptophan fluorescence with a second-order rate constant around (6.3±0.3)×10^3^ M^−1^s^−1^ for PfPrx6^WT^ or (2.0±0.4)×10^4^ M^−1^s^−1^ for hPrxVI (Fig. 1B-D, Table 1). The phases for PfPrx6^WT^ depended on the presence of the conserved histidyl residue and were lost for oxidized PfPrx6^H39Y^ (Fig. 1B). The results are consistent with the much slower DTT-dependent (*k* = 322 M^−1^s^−1^) reduction of oxidized PfPrx6, which also resulted in a decreased tryptophan fluorescence and required the presence of the histidyl residue.^12^ In contrast, we previously showed (and again confirmed, data not shown) that the formation of the sulfenic acid of PfPrx6^H39Y^ results in a change of tryptophan fluorescence similar to PfPrx6^WT^ and therefore does not require the histidyl residue.^8^ Thus, the negative result for the HS^−^-dependent reduction of oxidized PfPrx6^H39Y^ was not caused by altered tryptophan fluorescence properties. Furthermore, a negative control with reduced hPrxVI and Na_2_S revealed no reaction and therefore contradict the presence of other (contaminating) reactive sulfur species that might have oxidized the protein or altered the fluorescence in general (Fig. 1C). In summary, oxidized PfPrx6^WT^ and hPrxVI but not PfPrx6^H39Y^ rapidly react with HS^−^ as a potential universal electron donor for Prx6-type enzymes.

**Table 1.**
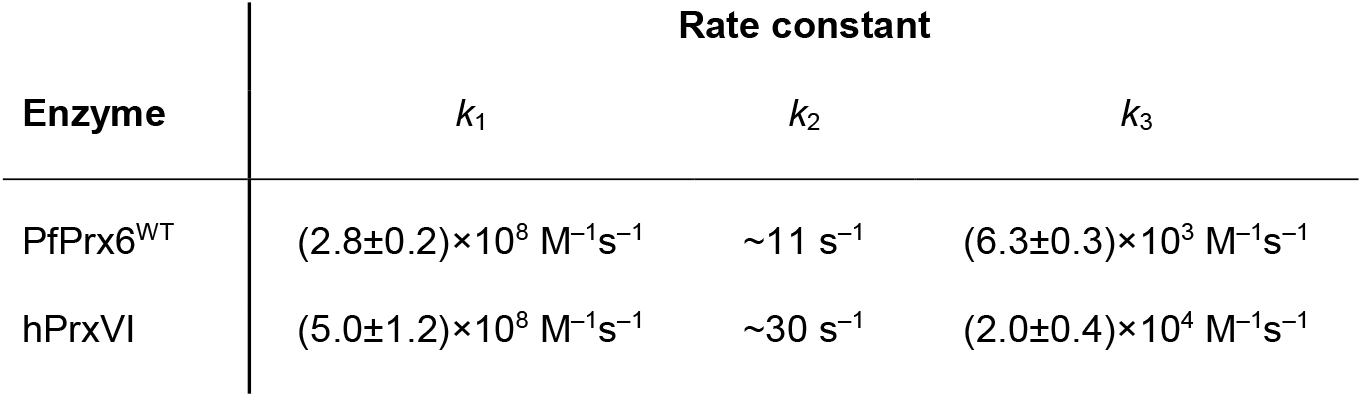
Rate constants from Figure 1 for the reactions of oxidized PfPrx6^WT^ or hPrxVI with Na_2_S at pH 7.4 and 25°C.

**Figure 1.**
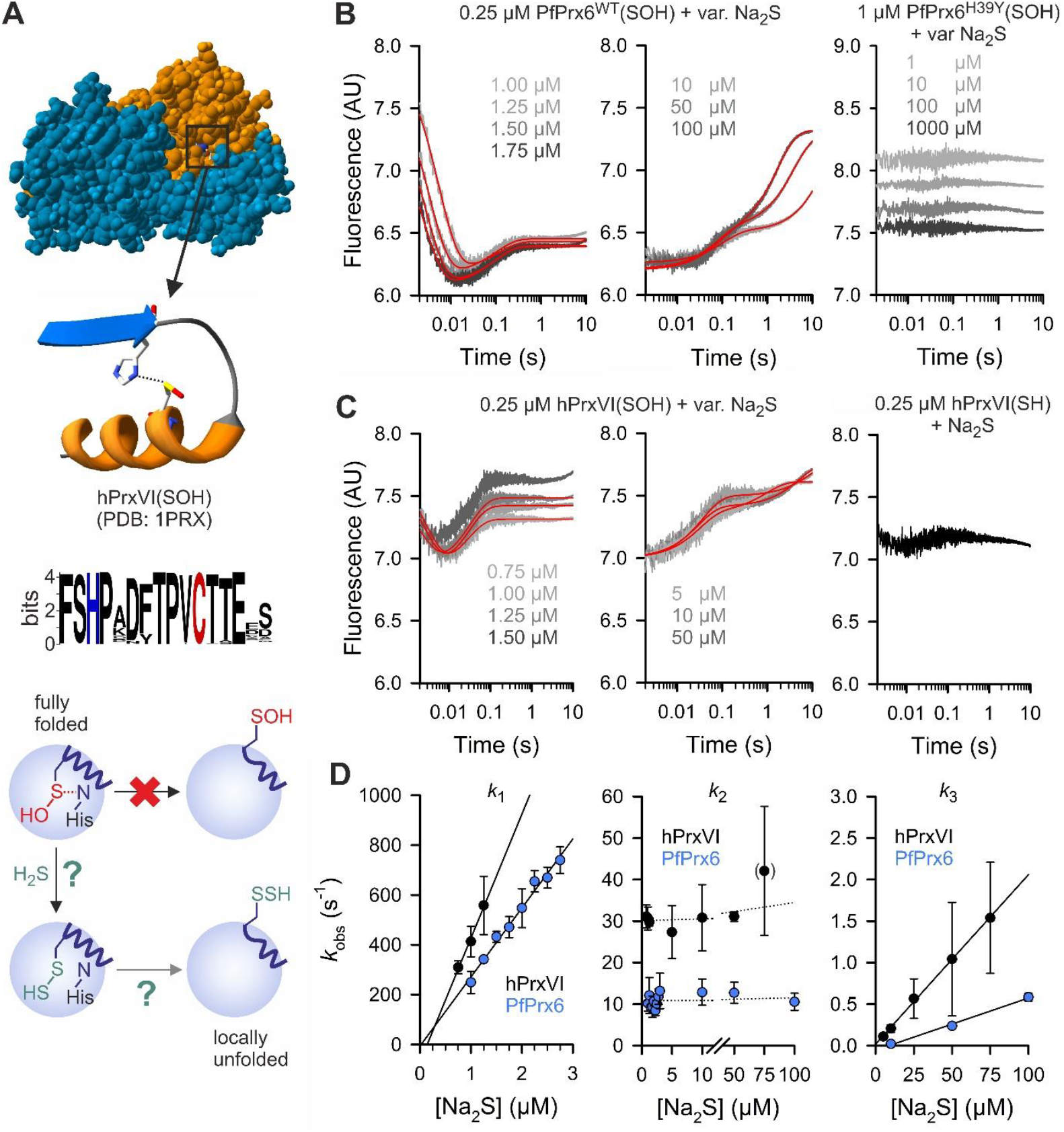
Rapid reduction of oxidized PfPrx6 by HS^−^. **A**) Structure of oxidized homodimeric hPrxVI (top), signature conservation graph for 5200 Prx6-type enzymes (middle),^11^ and schematic summary of a potential alternative HS^−^-dependent reduction mechanism of Prx6-type enzymes (bottom). The active-site cysteinyl and histidyl residue are highlighted for which the electron densities of crystallized hPrxVI and ApPrx6 were interpreted as a hydrogen bond or a hypervalent sulfur species, respectively.^20,21^ **B**) Representative kinetic traces of stopped-flow measurements with oxidized PfPrx6^WT^ (left panels) and PfPrx6^H39Y^ (right panel) at pH 7.4 and 25°C. Double exponential fits are shown in red. **C**) Representative kinetic traces of stopped-flow measurements with oxidized (left panels) and reduced hPrxVI (right panel) at pH 7.4 and 25°C. Double exponential fits are shown in red. **D**) Secondary plots of the *k*_obs_ values from the fits in panels B and C determined from three independent biological replicates. Rate constants were determined from the slope of the linear fits or the y-axis intercepts and are shown in Table 1.

### Working model for the HS^−^-dependent reduction of PfPrx6 and hPrxVI

While the exact chemical nature of the intermediates of the complex reduction pathway involving the active-site cysteinyl and histidyl residue remain to be characterized, we currently interpret the fluorescence changes for the HS^−^-dependent reduction of PfPrx6 and hPrxVI in Fig. 1B,C as follows: For the first reaction phase, we suggest a direct reaction between HS^−^ and the potentially hypervalent sulfur atom of the trapped oxidized species of both Prx6-type enzymes (Fig. 1A). Alternatively, following the elimination of water, HS^−^ could attack the sulfenamide between the active-site cysteinyl and histidyl residue. The resulting enzyme species of the first phase could be the protein persulfides PfPrx6^WT^(SSH) and hPrxVI(SSH) similar to the recently detected thioperselenide species Prx(SSeH) of hPrxVI that was formed with H_2_Se.^18,19^ Please note that analogous to the trapped, fully-folded sulfenic acid species of PfPrx6^WT^, which reacts with HS^−^ (Fig. 1B) but not cysteine,^12^ hPrxVI was shown to react with H_2_Se but not selenocysteine (which was explained by a steric inaccessibility of the sulfenic acid species).^19^ Also similar to the absent reaction of oxidized PfPrx6^H39Y^ with H_2_S (Fig. 1B), the ability of hPrxVI to transfer selenium to glutathione peroxidase 4 in SK-N-DZ cells depended on the presence of the active-site histidyl residue.^19^ In contrast to PfPrx6 and hPrxVI, the formation of a persulfide between the oxidized 1-Cys peroxiredoxin MtAhpE from *Mycobacterium tuberculosis* and Na_2_S was much slower with a second-order rate constant of (1.4±0.3)×10^3^ M^−1^s^−1^.^23^ A plausible explanation for the lower reactivity could be that a tyrosyl or phenylalanyl residue replaces the active-site histidyl residue in AhpE-type enzymes.^11^ The second phase in Fig. 1B,C might indicate an intramolecular rearrangement reflecting an equilibrium between a potential reaction product with a hypervalent sulfur atom and the potential free persulfide species PfPrx6^WT^(SSH) or hPrxVI(SSH) in the fully-folded protein conformation. The third phase might reflect the rather slow formation of fully reduced PfPrx6^WT^ or hPrxVI by excess Na_2_S. The final fluorescence after reaction with an excess of Na_2_S in Fig. 1B was slightly below the initial fluorescence of the oxidized enzyme, which is in accordance with a decreased fluorescence of the reduced enzyme as reported previously.^8^

### Excess HS^−^ fully reduces oxidized PfPrx6

To test our working model and to analyze if fully reduced PfPrx6^WT^ is generated in the presence of Na_2_S, we performed single-turnover experiments and additional controls (Fig. 2). For the single-turnover experiments, we either mixed H_2_O_2_ in the first syringe and reduced PfPrx6^WT^ with Na_2_S in the second syringe (Fig. 2A, left) or reduced PfPrx6^WT^ in the first syringe and freshly mixed H_2_O_2_ and Na_2_S in the second syringe (Fig. 2A, right). The alternative experimental designs were chosen to exclude a reaction between reduced PfPrx6^WT^ and potential polysulfides that might be present as impurities or that can be formed when H_2_O_2_ and Na_2_S are incubated for more than a few seconds. We observed similar changes of tryptophan fluorescence for both experimental designs, starting with a rapid loss of fluorescence (Fig. 2A), which probably reflects the formation of the sulfenic acid species of PfPrx6^WT^ as shown previously.^8^ While a further loss of fluorescence was detected after seconds with an equimolar concentration of Na_2_S in accordance with the loss of fluorescence in Fig. 1B, a concentration-dependent increase of fluorescence was observed with an excess of Na_2_S (Fig. 2A). We interpret this final phase in Fig. 2A as the formation of the potential persulfide species with one equivalent of HS^−^ (which presumably requires a rate-limiting conformational change or rearrangement after formation of the sulfenic acid species of PfPrx6^WT^)^8,12^, and the additional formation of the fully reduced enzyme by another equivalent of HS^−^ at higher Na_2_S concentrations. Similar changes in tryptophan fluorescence were obtained with an excess of DTT instead of Na_2_S (Fig. 2B), yielding the initial fluorescence at the end of a single-turnover experiment in accordance with the fully reduced enzyme. To show that the resulting enzyme species is indeed fully reduced PfPrx6^WT^, we analyzed its reactivity with H_2_O_2_ as a control (Fig. 2C). Pre-incubation of oxidized PfPrx6^WT^ with an excess of Na_2_S resulted in an enzyme species with identical kinetics with H_2_O_2_ as in ref. 8 when excess Na_2_S was removed (Fig. 2C, left). Here, the fluorescence increased beyond the initial fluorescence probably because of a conformational change or intramolecular rearrangement of the oxidized enzyme.^8^ In contrast, pre-incubation of oxidized PfPrx6^WT^ with an excess of Na_2_S resulted in an enzyme species with identical kinetics with H_2_O_2_ as in Fig. 2A when excess Na_2_S was present (Fig. 2C, right). Here, the initial and final fluorescence were identical in accordance with a reduced enzyme species before and after the reaction. In summary, while the intermediates of the complete reduction pathway remain to be characterized, our single-turnover experiments and controls support our working model and confirmed that the potential persulfide species of PfPrx6^WT^ can be reduced by an excess of HS^−^, yielding the fully reduced enzyme (Fig. 2D).

**Figure 2.**
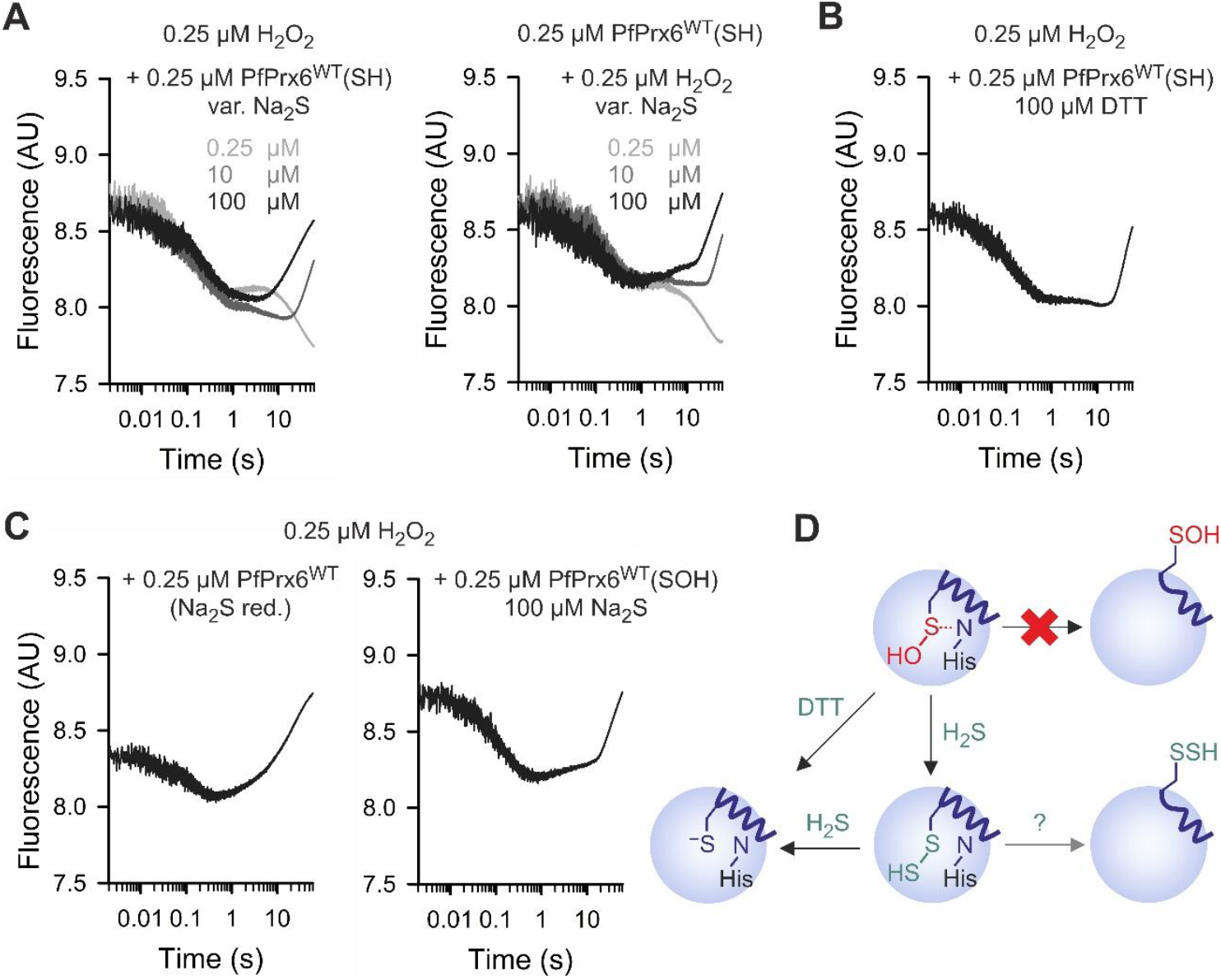
Excess HS^−^ fully reduces PfPrx6. **A**) Representative kinetic traces of single turnover stopped-flow measurements with either one equivalent of H_2_O_2_ in the first syringe and reduced PfPrx6^WT^ and the indicated concentrations of Na_2_S in the second syringe (left panel) or one equivalent of reduced PfPrx6^WT^ in the first syringe and freshly mixed H_2_O_2_ and the indicated concentrations of Na_2_S in the second syringe (right panel). **B**) Control experiment as in panel A with DTT replacing Na_2_S. **C**) Representative kinetic traces for the reaction between one equivalent of H_2_O_2_ in the first syringe and PfPrx6^WT^ that was oxidized and subsequently reduced with an excess of Na_2_S in the second syringe. Excess Na_2_S was either removed on a PD-10 column (left panel) or was still present in the reaction mixture (right panel). **D**) Schematic summary of the results. All concentrations indicate final concentrations in the mixing chamber. All experiments were performed at pH 7.4 and 25°C and all results were confirmed in an independent second biological replicate.

### The potential PfPrx6 persulfide does not react with the glutathione or thioredoxin systems

Next, we tested whether the potential (free or hypervalent) persulfide species of PfPrx6^WT^ reacts with alternative reducing agents including the glutathione and thioredoxin systems (Fig. 3).^13,24^ We therefore incubated oxidized PfPrx6^WT^ with one equivalent of Na_2_S and subsequently mixed the resulting enzyme species with either reduced thioredoxin (PfTrx1) or glutaredoxin (PfGrx) and GSH (Fig. 3A). Mutant PfGrx^C88S^ was used because it can react via a mono-or dithiol mechanism with glutathione disulfides and protein disulfides without potential side reactions involving a third cysteinyl residue.^25,26^ Since no change in fluorescence was detected, we excluded potential rapid reactions during the dead-time of the instrument by mixing oxidized PfPrx6^WT^ in the first syringe and Na_2_S and either reduced PfTrx1 or PfGrx^C88S^/GSH in the second syringe (Fig. 3B). No additional phases were observed as compared to Fig. 1B, indicating no reaction between potential PfPrx6^WT^(SSH) and PfTrx1 or PfGrx^C88S^/GSH. This is in contrast to the efficient reduction of several protein persulfides including PTP1B(SSH) and HSA(SSH) by Grx/GSH or Trx.^27-29^ We also tested the reaction between potential PfPrx6^WT^(SSH) and H_2_O_2_ or DTT (Fig. 3C). In contrast to reduced PfPrx6^WT^,^8^ no change in fluorescence occurred with H_2_O_2_ unless millimolar concentrations were used, resulting in a slow reaction. Complex changes in fluorescence were observed for the reaction of potential PfPrx6^WT^(SSH) with DTT. In contrast to the DTT-dependent reduction of PfPrx6^WT^(SOH),^12^ the reaction with potential PfPrx6^WT^(SSH) occurred much faster, and the number of phases varied depending on the DTT concentration used. The reactivity of potential PfPrx6(SSH), or the lack thereof, and the observed amplitudes overall suggest that the enzyme rests in the fully folded conformation. Likewise, incubation of hPrxVI with selenite and GSH was shown to yield exclusively the thioperselenide species PrxVI(SSeH) but no detectable adduct with glutathione.^18^ In summary, the reactivity of potential PfPrx6^WT^(SSH) is very different from reduced PfPrx6^WT^. Potential PfPrx6^WT^(SSH) can be reduced by Na_2_S or rapidly react in a complex mechanism with DTT, but neither accepts electrons from PfTrx1 or PfGrx^C88S^/GSH nor reacts with H_2_O_2_ as an oxidant (Fig. 3D).

**Figure 3.**
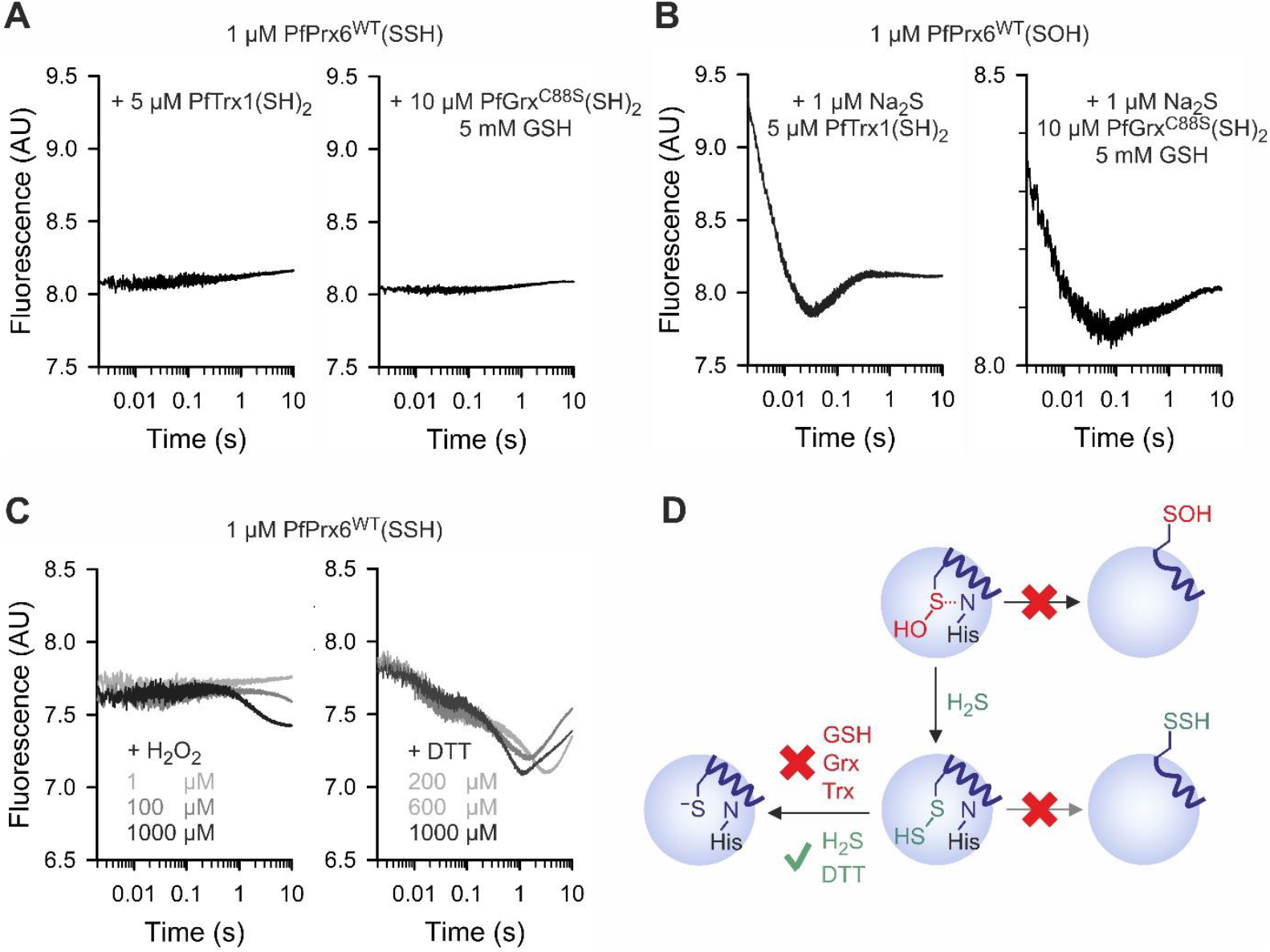
Potential PfPrx6(SSH) is not reduced by the thioredoxin or glutathione system. Representative kinetic traces of stopped-flow measurements with **A**) potential PfPrx6^WT^(SSH) or **B**) PfPrx6^WT^(SOH) and the indicated reductants. **C**) Representative kinetic traces for the reaction between potential PfPrx6^WT^(SSH) and H_2_O_2_ or DTT. **D**) Schematic summary of the results. All experiments were performed at pH 7.4 and 25°C and all results were confirmed in two independent biological replicates.

### Physiological implications of HS^−^ as a reducing agent for Prx6-type enzymes

While hPrxVI was recently shown to react with selenocysteine-derived H_2_Se with implications for ferroptosis,^18,19^ our data on PfPrx6 and hPrxVI suggest that HS^−^ is a potential universal reducing agent for Prx6-type peroxiredoxins. Unlike ascorbate, selenocysteine, or even GSH, H_2_S is present in all eukaryotes, bacteria, and archaea because of the fundamental biochemistry of iron-sulfur clusters. H_2_S can be an external (toxic) gas or serve as a signaling molecule or intracellular metabolite.^30,31^ A role of Prx6-type enzymes for H_2_S detoxification had been previously suggested based on high enzyme levels in lung tissue and epithelia as well as semi-quantitative *in vitro* data with bovine PrxVI.^3^ However, the 2-Cys Prx6-type enzyme from the annelid *Arenicola marina*, which lives in sediments that can contain high H_2_O_2_ and H_2_S concentrations, showed no activity in a steady-state assay with both redox reagents.^7^ Such negative results do not exclude a rapid reaction of the oxidized enzyme with HS^−^ and might be explained by (i) an enzyme inactivation due to hyperoxidation, (ii) a relatively slow second reduction of the protein persulfide by HS^−^, or (iii) the conversion of the sulfenic acid form of the studied 2-Cys Prx6-type enzyme via its persulfide species to its disulfide form without a net consumption of H_2_S. While we identified HS^−^ as a very efficient electron donor for the first reduction step of oxidized Prx6-type enzymes with a second-order rate constant >10^8^ M^−1^s^−1^, the second reduction by HS^−^ appears to be rather slow at physiological HS^−^ concentrations. The potential product of the second reduction is H_2_S_2_, which might react *in vivo* with millimolar GSH and serve as a source for glutathione persulfide (GSSH). GSSH could then react either with protein thiols or with another molecule of GSH, yielding GSSG. Such a Prx6-and HS^−^-dependent reduction of H_2_O_2_ could indeed explain the net formation of GSSG detected in yeast mitochondria.^15^ In summary, Prx6-type enzymes might either buffer fluctuations in H_2_S levels, act as H_2_S sensors that also integrate H_2_O_2_ signals, or require another partner for local unfolding to facilitate the reduction of the sulfenic acid or persulfide species as shown for oxidized hPrxVI and glutathione transferase P1-1 in ref. 14. Follow-up studies are necessary to characterize the intermediates and the exact reaction mechanism for the HS^−^-dependent reduction of PfPrx6, hPrxVI, and other Prx6-type enzymes, and to study the physiological relevance of Prx6-type enzymes for the detoxification of external H_2_S, for H_2_S-dependent signaling, and for iron-and sulfur metabolism.

### Experimental procedures

#### Materials

Sodium sulfide (Na_2_S), dithiothreitol (DTT), and reduced L-glutathione (GSH) were from Sigma Aldrich, diethylenetriaminepentaacetic acid (DTPA) was from Carl Roth, H_2_O_2_ from VWR, and isopropyl-*β*-D-1-thiogalactopyranoside (IPTG) was from Serva.

#### Molecular biology, heterologous expression, and protein purification

The gene encoding hPrxVI was PCR-amplified with Phusion HF DNA polymerase using forward primer GATC*<u>CCATGG</u>*GACCCGGAGGTCTGCTTCTCG, reverse primer GATC*<u>CTCGAG</u>*AGGCTGGGGTGTGTAGCGG, and plasmid p413/*HSPRDX6-FLAG* as a template^15^. The product was subcloned into the *Nco*I and *Xho*I restriction sites of pET28a/*PFPRX6-His6* in *Escherichia coli* strain XL1-Blue, yielding pET28a/*HSPRXVI-His6*. The correct sequence was confirmed by Sanger sequencing of both strands (Microsynth Seqlab). Recombinant C-terminally LEH_6_-tagged PfPrx6^WT^, PfPrx6^H39Y^, and hPrxVI were produced in *E. coli* strain SHuffle T7 express (NEB) at 30°C and N-terminally MRGSH_6_GS-tagged PfGrx^C88S^ and PfTrx1 were produced in *E. coli* strain XL1-Blue at 37°C. Gene expression was induced at an OD_600nm_ of 0.5 with 0.5 mM IPTG for 4 hours. *E. coli* cultures were then cooled in an ice-water bath for 15 minutes and centrifuged (15 min, 4°C, 3750×g). Cell pellets were resuspended in ice-cold buffer I (20 mM imidazole, 100 mM Na_x_H_y_PO_4_, 300 mM NaCl, pH 8.0 at 4°C). Cell suspensions were stirred on ice with DNaseI and 10 mg lysozyme for 45 minutes and then sonicated. The cell lysates were centrifuged (30 min, 4°C, 10000×g) and the supernatants loaded on pre-equilibrated Ni-NTA agarose (Qiagen) columns. The columns were washed with 15 column volumes of ice-cold buffer I and proteins were eluted with ice-cold buffer II (200 mM imidazole, 100 mM Na_x_H_y_PO_4_, 300 mM NaCl, pH 8.0 at 4°C). The purity of the eluates was confirmed by analytical SDS-PAGE. Activities of PfPrx6^WT^, PfPrx6^H39Y^, and hPrxVI were confirmed with H_2_O_2_, the activity of PfTrx1 was confirmed with 5,5′-dithiobis-(2-nitrobenzoic acid), and the activity of PfGrx^C88S^ was confirmed with bis(2-hydroxyethyl)disulfide as described previously.^8,12,32^

#### Sample preparation

Freshly purified enzymes were reduced with 5 mM DTT for 30 minutes on ice. Excess DTT and imidazole were removed on a PD-10 desalting column (Merck) and reduced proteins were eluted with 3.5 ml ice-cold assay buffer (100 mM Na_x_H_y_PO_4_, 0.1 mM DTPA, pH 7.4 at 25°C). Protein concentrations were determined spectrophotometrically at 280 nm using extinction coefficients of 30940 M^−1^cm^−1^, 32430 M^−1^cm^−1^, or 22460 M^−1^cm^−1^ for reduced PfPrx6^WT^, PfPrx6^H39Y^, or hPrxVI, respectively. Oxidized PfPrx6^WT^, PfPrx6^H39Y^, or hPrxVI were generated by incubation of the reduced enzymes with equimolar H_2_O_2_ for 30 min on ice. The potential persulfide species of PfPrx6^WT^ and PfPrx6^H39Y^ were generated by incubation of the oxidized enzyme species with equimolar Na_2_S for 15 min on ice.

#### Stopped-flow kinetic measurements

Kinetic measurements were carried out in a thermostatted SX-20 spectrofluorometer (Applied Photophysics) at 25°C. The change of tryptophan fluorescence was measured as total emission at an excitation wavelength of 295 nm with a slit width of 2 mm. All reagents were freshly dissolved in ice-cold assay buffer. For the reaction of oxidized PfPrx6^WT^, PfPrx6^H39Y^, or hPrxVI with Na_2_S, 0.5 μM freshly H_2_O_2_-oxidized enzyme in syringe 1 was mixed with different concentrations of excess Na_2_S in syringe 2. Traces of at least three technical replicates were averaged and fitted by double exponential regression using the Pro-Data SX software (Applied Photophysics) to obtain *k*_obs_ values. The *k*_obs_ values of three biological replicates were plotted against the Na_2_S concentration in SigmaPlot 13.0 to obtain rate constants for each phase from linear fits. Single-turnover experiments were performed with either one equivalent of H_2_O_2_ in the first syringe and reduced PfPrx6^WT^ and variable concentrations of Na_2_S in the second syringe or one equivalent of reduced PfPrx6^WT^ in the first syringe and freshly mixed H_2_O_2_ and Na_2_S (with an incubation time well under one minute) in the second syringe. The influence of the Trx or Grx/GSH systems was monitored by mixing 2 μM freshly H_2_O_2_-oxidized PfPrx6^WT^ in syringe 1 with either 2 μM Na_2_S and up to 50 μM PfTrx1 or 2 μM Na_2_S, 10 mM GSH, and up to 50 μM PfGrx^C88S^ in syringe 2. Reactions of the potential PfPrx6 persulfide were investigated by mixing 2 μM freshly Na_2_S-incubated PfPrx6^WT^(SOH) in syringe 1 with up to 50 μM PfTrx1, up to 50 μM PfGrx^C88S^ with 10 mM GSH, up to 2 mM DTT, or up to 2 mM H_2_O_2_ in syringe 2.

## Data availability

Data described in the manuscript are contained within it. Additional raw data is available upon request to Marcel Deponte at.

## Acknowledgements

This work was funded by the Deutsche Forschungsgemeinschaft (DFG) grant DE 1431/19-1 to M.D. (project number 508372800). La.Le., L.T., and M.D. are supported by the DFG RTG 2737 (STRESSistance). We thank Jan Riemer for plasmid p413/*HSPRDX6-FLAG*.

## Author Contributions

Lu.La., La.Le., and L.T. performed the experiments and analyzed the data. M.D. and Lu.La. conceptualized the study. M.D. supervised the study. M.D. and Lu.La. wrote the manuscript. All authors read and approved the final version of the manuscript.

## Competing Interests

The authors declare no competing interests.

### Abbreviations

DTPA: diethylenetriaminepentaacetic acid
DTT: dithiothreitol
IPTG: isopropyl-*β*-D-1-thiogalactopyranoside
GSH: reduced glutathione
Grx: glutaredoxin(s)
Trx: thioredoxin(s)

